# Wnt/β-catenin signaling controls mouse eyelid growth by mediating epithelial-mesenchymal interactions

**DOI:** 10.1101/2023.05.14.540693

**Authors:** Xuming Zhu, Makoto Senoo, Sarah E. Millar, Gang Ma

**Author notes:** Author for correspondence: Sarah E Millar Gang Ma.

## Abstract

Eyelid closure is required for the development of multiple ocular tissues. Interactions between the epithelium and underlying mesenchyme play a pivotal role in regulating eyelid development. However, the molecular mechanisms underlying these interactions remain unclear. Wnt signaling pathways regulate the development of ocular tissues, but their functions in eyelid development are not fully defined. In this study, we find that deletion of β-catenin in supraorbital mesenchyme abolishes eyelid growth by causing decreased proliferation in supraorbital epithelium and underlying mesenchyme. Inhibition of Wnt secretion by deleting *Wls* in supraorbital epithelium results in failure of eyelid developing, similar to the effects of deleting mesenchymal β-catenin, suggesting that mesenchymal Wnt/β- catenin signaling is controlled by epithelial Wnt ligands during eyelid development. We also found that deletion of *p63* results in formation of hypoplastic eyelids and reduced expression of several Wnt ligands in eyelid epithelium, indicating that expression of Wnt ligands in eyelid development is at least partially regulated by p63. Taken together, our data indicate that Wnt/β-catenin signaling controls eyelid growth by orchestrating epithelial- mesenchymal interactions.

## Introduction

Eyelid colobomas, characterized by partial or complete loss of eyelid tissue, occur in more than twenty human syndromes [1]. However, the mechanisms controlling eyelid development, and how these are perturbed to cause defective eyelid development, are incompletely understood [2].

The mammalian eyelid originates from periocular epithelium and underlying mesenchyme, and its development can be categorized into four main stages comprising specification, growth, fusion, and separation [3, 4]. Our current knowledge of the mechanisms of eyelid development is largely based on murine genetic studies, as a diverse collection of genetically modified mouse mutants displays an eye-open at birth (EOB) phenotype [4]. The course of mouse eyelid formation closely resembles that of human eyelid, making it an ideal model to study eyelid development.

Murine eyelid development initiates at approximately embryonic day 9 (E9) when the presumptive eyelid region can be identified by *Pax6* expression [5]. By E11.5, invagination of both dorsal and ventral periocular tissue outlines morphologically discernable eyelids, which continue to grow toward each other across the ocular globe. Starting at E15.5, the peridermal cell layers of both eyelids extend from the leading edges and fuse across the cornea by E16.5. Thereafter, the palpebral epidermis that bridges the eyelids begins to stratify, and other epidermal appendages such as hair follicles and Meibomian glands begin to develop. After birth, the eyelids are fused and subsequently start to separate. Separation of eyelids, and opening of eyes, is complete by postnatal day 12 (P12) [6].

EOB phenotypes in most of the mouse mutants that have been characterized to date are caused by impaired eyelid closure, and are associated with altered expression and/or activity of a diverse set of transcription factors and signaling pathways. For example, Activin βB regulates eyelid closure by triggering a signaling cascade that integrates MAP3K1, JNK and c-JUN activities [7, 8]. c-JUN may also induce EGFR expression, which mediates the effect of EGF in promoting periderm migration via activation of the ERK pathway [9, 10]. The FGF10-FGFR2 pathway is required for eyelid epithelial proliferation and promotes periderm migration by inducing expression of Activin βB and TGF-α [11]. Additionally, FGF signaling acts upstream of the SMAD4-mediated BMP pathway in maintaining FOXC1 and FOXC2 expression, which regulate eyelid closure in a dosage-dependent manner [12, 13]. While the regulation of eyelid fusion has been intensively investigated, the mechanisms regulating eyelid growth are less well characterized.

Wnt proteins are secreted paracrine factors that trigger both β-catenin-dependent canonical and β-catenin-independent non-canonical Wnt pathways. The cargo protein WNTLESS (WLS) is required for secretion of Wnt ligands [14, 15]. Multiple Wnts are expressed in the periocular tissue during eyelid development [16], and Wnt/β-catenin signaling is reported to regulate eyelid closure by repressing MAP3K1 expression and JNK activity [17]. Deletion of DKK2, a sectreted Wnt inhibitor, also causes abnormal eyelid development, indicating that excessive Wnt activity affects eyelid closure [18]. Therefore, the Wnt signaling pathway must be precisely controlled for normal eyelid development.

In the current study, we investigated the functions of Wnt/β-catenin signaling in eyelid growth by genetically manipulating the expression of mesenchymal β-catenin and epithelial *Wls*. We found that deletion of either mesenchymal β- catenin or epithelial *Wls* disrupts eyelid growth. In addition, we observed that expression of Wnt ligands in eyelid epithelium is decreased in *p63* loss of function mutants, which have hypoplastic eyelids, indicating that epithelial Wnt expression is at least partially regulated by p63 during eyelid development.

## Materials and methods

### Mice

The following mouse lines were used: *Prx1-Cre* (Jackson Laboratories, strain #005584); *Msx2-Cre* (Jackson Laboratories, strain #027892); *ROSA^mT/mG^* (Jackson Laboratories, strain #007676); *ROSA26^LacZ^* (Jackson Laboratories, strain #002073); *Wls^fl/fl^* (Jackson Laboratories, strain #012888); β*-catenin^fl/fl^* (Jackson Laboratories, strain #004152); and *p63^+/-^*[19]. All mice were maintained on a mixed strain background. Mice were allocated to experimental or control groups according to their genotypes, with control mice included in each experiment. Male mice carrying *Prx1-Cre* or *Msx2-Cre* were crossed with *ROSA^mT/mG^*, *ROSA26^LacZ^*, β*-catenin^fl/fl^*, or *Wls^fl/fl^* females to avoid potential deletion in the female germline. *p63^+/-^* male and female mice were intercrossed to yield *p63^-/-^* embryos. Investigators were not blinded during allocation and animal handling as information about genotype was required for appropriate allocation to experimental groups. Immunostaining and RNA in situ hybridization studies were carried out and results recorded in a blinded fashion. Up to five mice were maintained per cage in a specific pathogen-free barrier facility on standard rodent laboratory chow (Purina, catalog #5001). All animal experiments were performed under approved animal protocols according to institutional guidelines established by the respective IACUC committees at the Icahn School of Medicine at Mount Sinai and Shanghai Jiaotong University.

### RNA in situ Hybridization (ISH)

Embryos at the indicated stages were harvested, fixed with 4% paraformaldehyde (PFA) (Affymetrix/USB) in PBS overnight at 4°C, and processed for whole-mount RNA ISH, or frozen or paraffin sectioning. Whole-mount RNA ISH procedures and the *Fgf10* ISH probe were previously described [20]. RNAscope was performed on fixed frozen sections or paraffin sections following the user’s guide provided by Advanced Cell Diagnostic (ACD) using probes for *Lef1* (ACD #441861), *Wnt4* (ACD #401101), *Wnt3* (ACD #312241), and *Twist1* (ACD #414701).

### ***β***-Galactosidase staining

Embryos at the indicated stages were harvested and fixed with 4% paraformaldehyde (PFA) (Affymetrix/USB) in PBS for 30 minutes at room temperature. Subsequent procedures were carried out as previously described [21].

### BrdU administration

20mg/ml BrdU (B5002, Sigma) in PBS was used for BrdU labeling. Pregnant female mice were intraperitoneally injected with 2mg BrdU per 30 g body weight. After 1 hour, the mice were euthanized to collect embryos at the indicated stages.

### Immunostaining and TUNEL assays

Frozen sections were incubated with Ki-67 antibody (#14-5698-82, Invitrogen, 1:200), β-catenin antibody (#C7207, Sigma, 1:1000), BrdU antibody (#B2531, Sigma, 1:200), LEF1 antibody (#2230s, Cell Signaling Technology), KRT5 antibody (#905501, BioLegend, 1:500) and KRT14 antibody (#MA5-11599, Invitrogen, 1:500) overnight at 4°C, followed by incubation with Alexa Fluor-labelled secondary antibodies (Thermo Fisher Scientific). For quantification of proliferating cells, at least 30 epithelial and 100 underlying mesenchymal cells in the presumptive upper eyelid from each embryo were analyzed. Unpaired two-tailed Student’s *t*-test was used to calculate statistical significance for quantification of Ki-67+ or BrdU+ cells, and p<0.05 was considered significant. For TUNEL assays, fixed frozen sections were incubated with TUNEL labeling mix (Roche) at 37°C for 1 hour and then washed with PBST and mounted in DAPI containing medium (Thermo Fisher Scientific). All immunofluorescent (IF) data were documented using a Leica Microsystems DM5500B fluorescent microscope (Leica Microsystems).

## Results

### *Prx1-Cre* is active in supraorbital and developing upper eyelid mesenchyme

To determine the role of canonical Wnt pathway during eyelid growth, we sought to delete β-catenin in eyelid mesenchyme which displays a high level of Wnt activity as evidenced by expression of the ubiquitous Wnt target gene, *Axin2* [22]. To identify a Cre line that is active in eyelid mesenchyme, we first examined the activity of *Prx1-Cre* in periocular tissue during eyelid development because *Prx1-Cre* was reported to be active in a subset of cranial-facial mesenchymal cells [23]. *Prx1-Cre* mice were crossed with the *Rosa26R^mT/mG^* Cre reporter allele to monitor Cre activity. We found that at E11.5, reporter gene activity was present in a subset of supraorbital mesenchyme of the presumptive upper eyelid (Fig. S1B-D). By E12.5, Cre reporter expression in the supraorbital mesenchyme had expanded rostrally, and increased numbers of deleted cells were detected in the developing upper eyelid primordia (Fig. S1E-G). By E13.5, the domain of reporter expression continued to expand, and the entire upper eyelid and surrounding mesenchyme were positive for mGFP expression (Fig. S1H-J). These data suggest that *Prx1-Cre* can be used to manipulate gene expression in supraorbital and upper eyelid mesenchyme.

### Deletion of **β**-catenin in supraorbital mesenchyme abolishes upper eyelid growth

To test whether mesenchymal canonical Wnt signaling regulates eyelid development, we used *Prx1-Cre* to delete β-catenin. We found that eyelid development at E18.5 was unaffected by heterozygous deletion of β-catenin in *Prx1-Cre* β*-catenin^fll+^* embryos compared with wild-type. However, homozygous deletion in *Prx1-Cre* β*-catenin^fl/fl^*embryos resulted in an open eye phenotype (Fig. 1A and 1B). H&E staining of sectioned tissue revealed that the upper eyelids of these mutants were almost completely absent except for some rudimentary epidermal thickening, while the lower eyelids still developed (Fig. 1C and 1D). This observation was consistent with differential *Prx1-Cre* activity between upper and lower eyelid mesenchyme (Fig. S1). To exclude the possibility that the upper eyelid defect was caused by abnormal eyelid fusion, we examined eyelids at E12.5 and E14.5 when eyelid fusion has not initiated. We found that the upper eyelids of *Prx1-Cre* β*-catenin^fl/fl^*embryos failed to develop at E12.5 and E14.5 (Fig. 1E-H). These data indicate that supraorbital mesenchymal β-catenin is required for upper eyelid growth.

**Figure 1.**
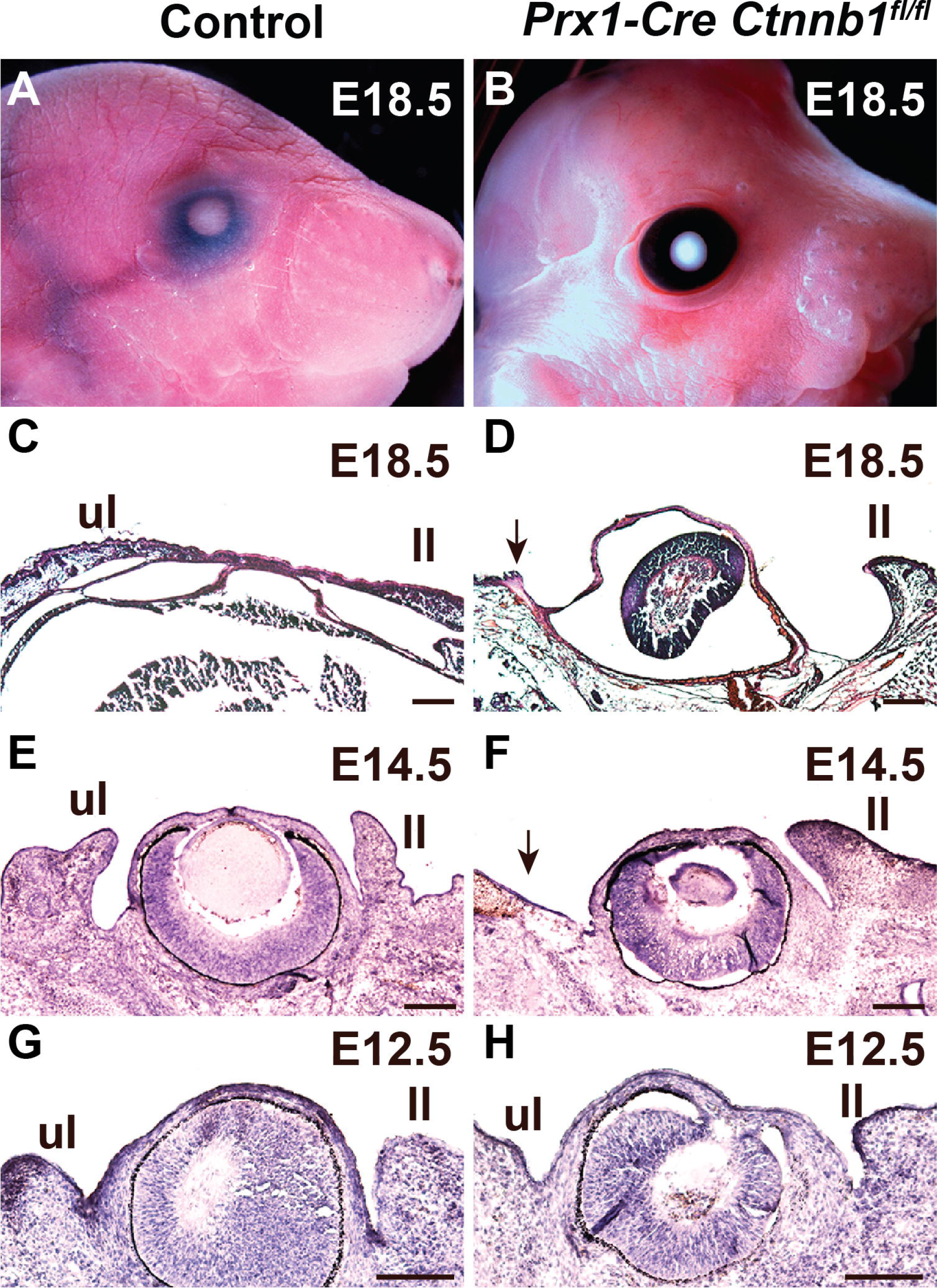
Deletion of β-catenin in supraorbital mesenchyme abolishes upper eyelid growth. (A) The eyelids of control embryos are fused at E18.5. (B) The eyes of *Prx1-Cre* β*-catenin^fl/fl^* embryos are open at E18.5. (C) H&E staining of tissue sections shows that eyelids are fused in control embryos at E18.5. (D) H&E staining of tissue sections indicates that the upper eyelid is almost absent in *Prx1-Cre* β*-catenin^fl/fl^* embryos, except for some epidermal thickening (black arrow). The lower eyelid is still present in these mutants. (E) H&E staining of tissue sections shows that control eyelids are well-formed at E14.5. (F) H&E staining of tissue sections indicates that there is no discernable upper eyelid in *Prx1-Cre* β*-catenin^fl/fl^*embryos at E14.5 (black arrow). (G) H&E staining shows that at E12.5, control embryos have morphologically distinguishable eyelid primordia. (H) H&E staining shows that in *Prx1-Cre* β*-catenin^fl/fl^*embryos, no significant upper eyelid primordia have formed at E12.5, but lower eyelid primordia are present. Scale bars: 150μm. Abbreviations: ul, upper eyelid; ll, lower eyelid.

### Wnt/**β**-catenin signaling in supraorbital mesenchyme regulates cell proliferation

To delineate the mechanisms by which β-catenin regulates eyelid growth, we assessed activity of the canonical Wnt pathway in the supraorbital region when eyelid growth initiates at E11.5. We found that supraorbital mesenchymal β-catenin was efficiently deleted by *Prx1-Cre* at E11.5, although small patches of undeleted cells could still be identified (Fig. 2A and 2B). Expression of *Lef1* and *Twist1*, which are direct Wnt/β-catenin target genes [24], was significantly reduced in β-catenin-deficient supraorbital mesenchyme (Fig. 2C-F). By contrast, expression of epithelial Wnt ligands including *Wnt3* and *Wnt4* was not affected in these mutants (Fig. 3G-J). To determine whether increased apoptosis contributes to the eyelid defects in *Prx1-Cre* β*-catenin^fl/fl^* embryos, we performed TUNEL assays. The results indicated absence of abnormal apoptosis in the supraorbital region of the mutants at E11.5 (Fig. 3K and 3L). Since Wnt/β-catenin pathway regulates proliferation of mesenchymal cells in multiple contexts [20, 25], we asked whether proliferation was affected in the supraorbital region of *Prx1-Cre* β*-catenin^fl/fl^*mutants at E11.5. The data indicated that proliferation was decreased in both supraorbital mesenchyme and overlying epithelium in the mutant eyelids compared with controls (Fig. 3M-O). Taken together, these data suggest that mesenchymal β-catenin regulates cell proliferation in the presumptive upper eyelid.

**Figure 2.**
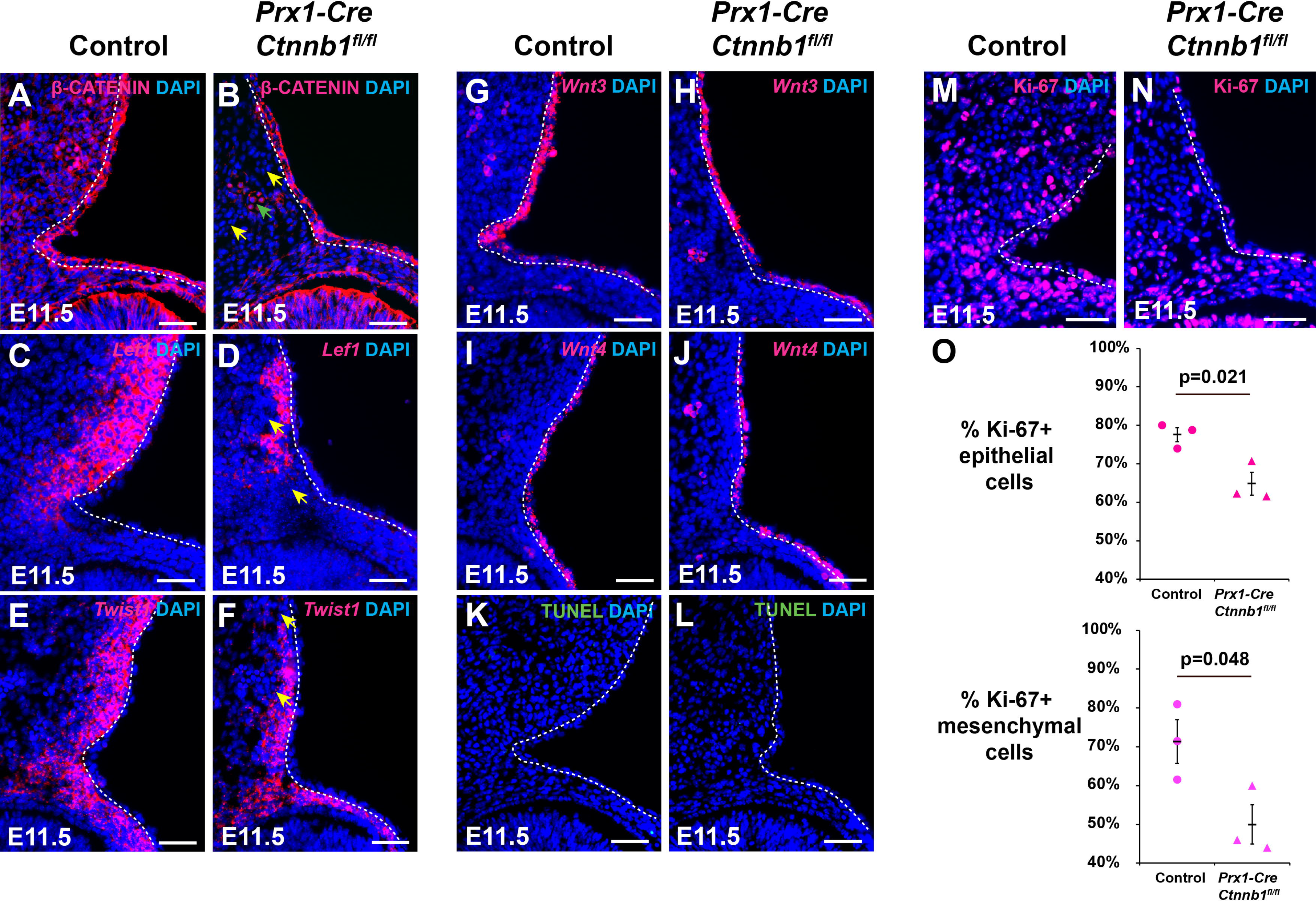
Deletion of. β**-catenin in supraorbital mesenchyme results in reduced canonical Wnt activity and decreased cell proliferation.** (A) IF shows that β-catenin is expressed in the supraorbital epithelium and mesenchyme of control embryos at E11.5. (B) IF shows that β-catenin is deleted in the supraorbital mesenchyme (yellow arrows), but patches of undeleted cells can also be identified (green arrow) in *Prx1-Cre* β*-catenin^fl/fl^* embryos at E11.5. (C) *Lef1* expression is detected in supraorbital mesenchyme in control embryos at E11.5. (D) *Lef1* expression is reduced or absent (yellow arrows) in the supraorbital mesenchyme of *Prx1-Cre* β*- catenin^fl/fl^*embryos at E11.5. (E) *Twist1* is expressed in the supraorbital mesenchyme of control embryos at E11.5. (F) Expression of *Twist1* is decreased in the supraorbital mesenchyme of *Prx1-Cre* β*-catenin^fl/fl^* embryos at E11.5 (yellow arrows). (G and H) Expression of epithelial *Wnt3* is similar between control and *Prx1-Cre* β*-catenin^fl/fl^* embryos at E11.5. (I and J) Epithelial *Wnt4* levels are comparable between control and *Prx1-Cre* β*- catenin^fl/fl^*embryos at E11.5. (K and L) TUNEL assay indicates that levels of apoptosis are similar in control and *Prx1-Cre* β*-catenin^fl/fl^*embryos at E11.5. (M and N) Ki-67 staining at E11.5 shows that proliferation is reduced upon deletion of mesenchymal β-catenin. (O) Statistical analysis of Ki-67+ cells in supraorbital epithelium and underlying mesenchyme. Three pairs of control and mutant embryos were analyzed. Unpaired student *t*-test was used to evaluate significance. p<0.05 was considered significant. Scale bars: 50μm. The white dashed line indicates the boundary between epithelium and mesenchyme.

**Figure 3.**
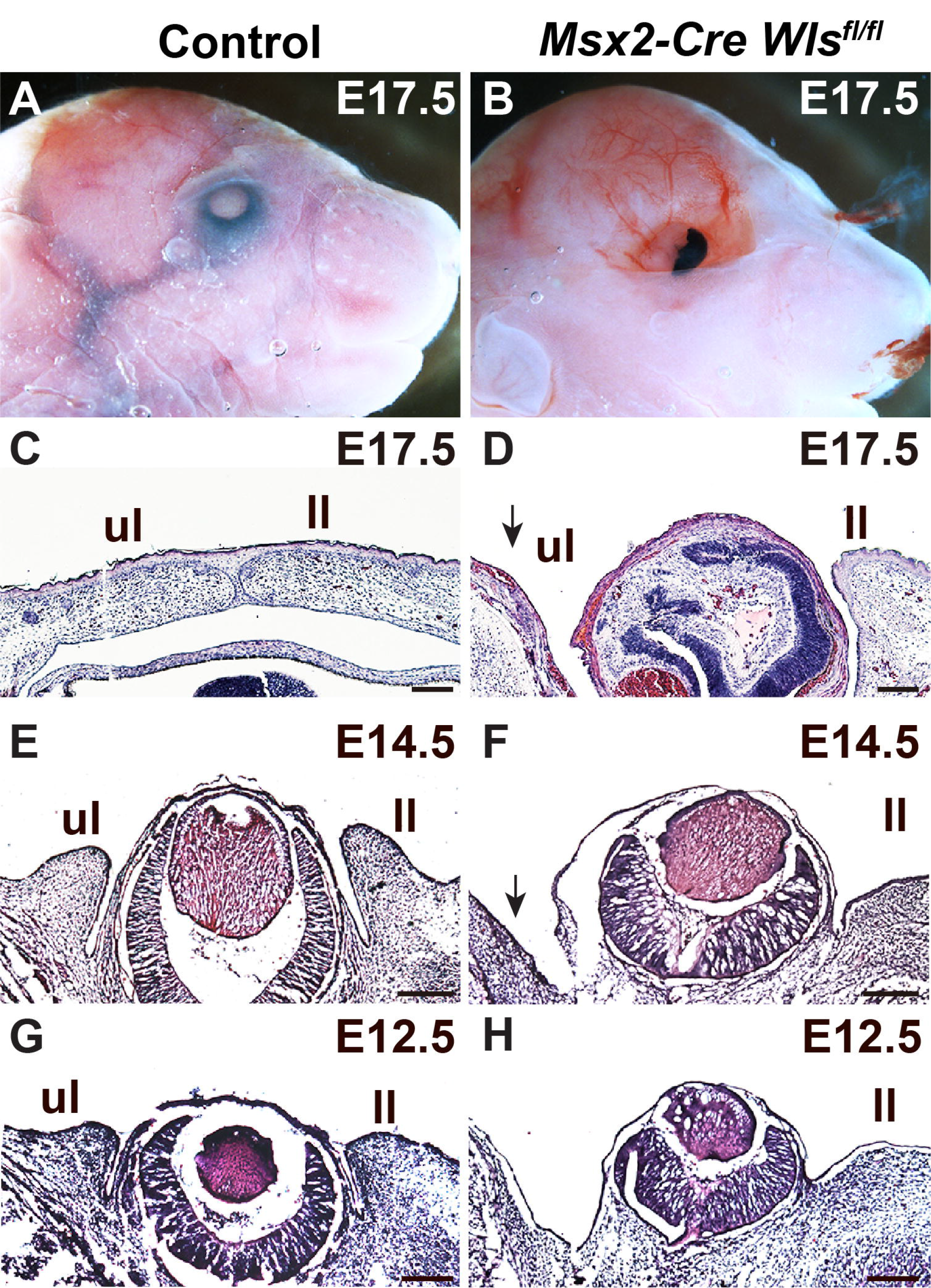
Epithelial Wnt ligands are required to maintain eyelid growth. (A) The eyelids of control embryos are fused at E17.5. (B) The eyes of *Msx2- Cre Wls^fl/fl^* embryos exhibit defects in multiple ocular tissues including loss of supraorbital skin and eyelid, and malformed eye at E17.5. (C) H&E staining shows that eyelids are fused in control embryos at E17.5. (D) H&E staining indicates that the upper eyelid is lost in *Msx2-Cre Wls^fl/fl^* embryos (black arrow). The lower eyelid is still present in these mutants. (E) H&E staining shows that control eyelids are well-formed at E14.5. (F) H&E staining indicates that the upper eyelid is lost in *Msx2-Cre Wls^fl/fl^* embryos at E14.5 (black arrow). (G) H&E staining shows that at E12.5, control embryos have morphologically distinguishable eyelid primordia. (H) H&E staining shows that in *Msx2-Cre Wls^fl/fl^* embryos, no upper eyelid primordia have formed at E12.5. Scale bars: 150μm. Abbreviations: ul, upper eyelid; ll, lower eyelid.

### Epithelial Wnt ligands are required to maintain eyelid growth

Since Wnt genes are predominantly expressed in eyelid epithelium [16], we investigated whether epithelial Wnt ligands are necessary for eyelid growth. To disrupt the function of epithelial Wnt ligands, we blocked their secretion y deleting *Wls* with *Msx2-Cre*, which is active in periocular epithelium, including the presumptive eyelid epithelium at E11.5 and the developing eyelid epithelium from E12.5 to E13.5 (Fig. S2B-J). We found that *Msx2-Cre Wls^fl/fl^* embryos exhibited multiple defects in ocular tissue development including absence of normal eyelids at E17.5 (Fig. 3A and 3B). H&E staining data showed that the upper eyelids failed to form, and the lower eyelids were hypoplastic (Fig. 3C and 3D) in these mutants. Examination of eye tissue at E12.5 and E14.5 confirmed that growth of upper was completely disrupted in the mutants (Fig. 3E-H). These data demonstrate that epithelial Wnt ligands are necessary for eyelid growth.

### Epithelial Wnt ligands control supraorbital mesenchymal canonical Wnt signaling

Since *Msx2-Cre Wls^fl/fl^* mutants displayed similar defects in eyelid development to those observed in *Prx1-Cre* β*-catenin^fl/fl^*mutants, we asked whether epithelial Wnt ligand secretion is required for canonical Wnt signaling in the mesenchyme. We found that epithelial *Wls* was efficiently deleted in *Msx2-Cre Wls^fl/fl^* mutants (Fig. 4A and 4B), and expression of the Wnt/β- catenin target genes LEF1 and *Twist1* was downregulated in the underlying mesenchyme, indicating reduced canonical Wnt activity (Fig. 4C-F). By contrast, mRNA levels of *Wnt3* and *Wnt4* were comparable between control and *Wls*-deficient epithelium (Fig. 4G-J), indicating that *Wls* deletion blocked Wnt protein secretion but did not affect expression of Wnt genes. *Msx2-Cre Wls^fl/fl^* mutants displayed reduced proliferation in both supraorbital epithelium and mesenchyme, but increased apoptosis was not observed (Fig. 4K-O). These observations demonstrate that epithelial Wnts regulate eyelid growth by activating canonical Wnt signaling in supraorbital mesenchyme.

**Figure 4.**
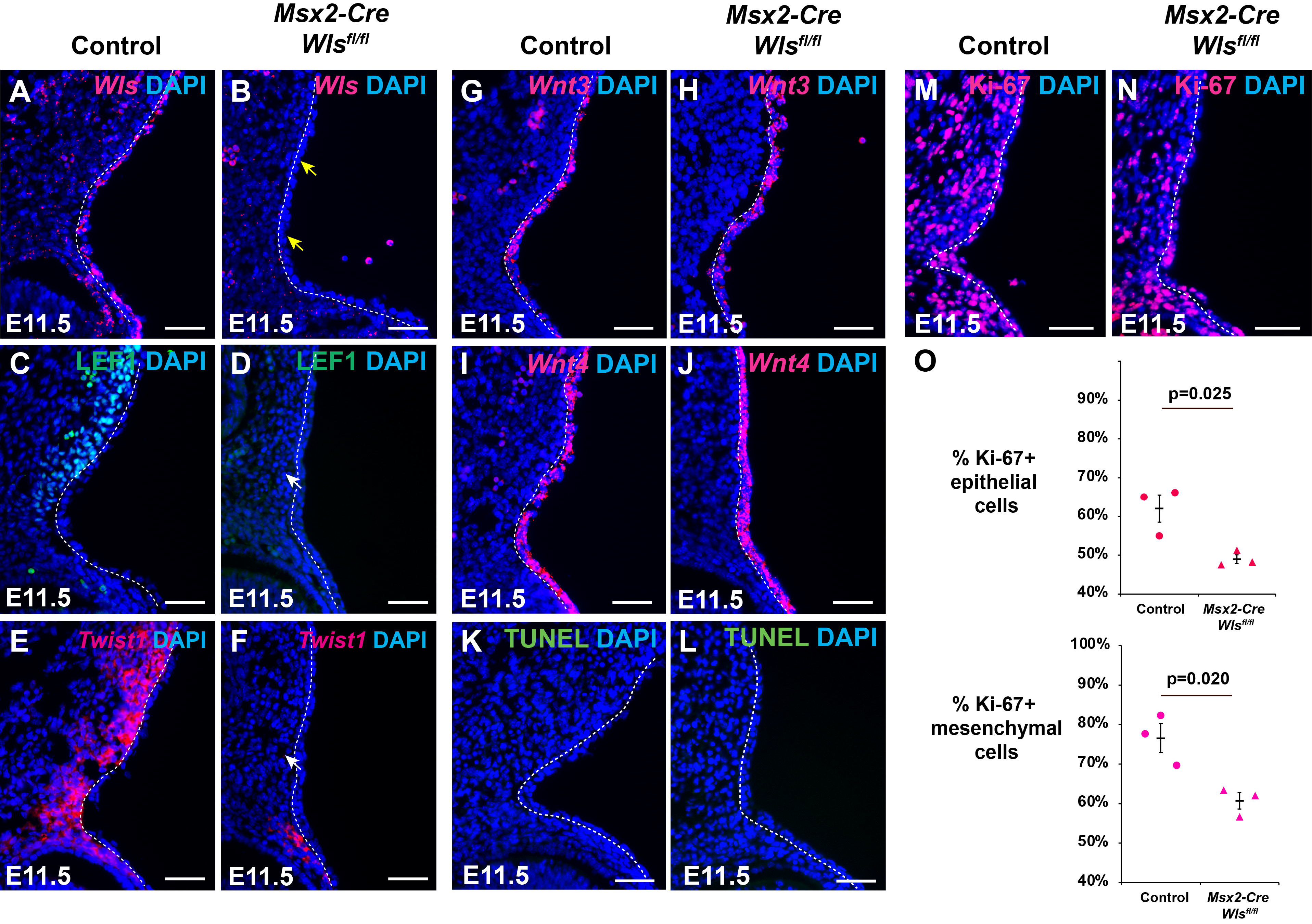
Epithelial Wnt ligands control mesenchymal canonical Wnt signaling in the eyelid. (A) RNAscope ISH shows that *Wls* is expressed in the supraorbital epithelium and mesenchyme of control embryos at E11.5. (B) RNAscope ISH shows that *Wls* is deleted in the supraorbital epithelium (yellow arrows) of *Msx2-Cre Wls^fl/fl^* embryos at E11.5. (C) IF staining shows that LEF1 is expressed in the supraorbital mesenchyme of control embryos at E11.5. (D) IF staining shows that LEF1 expression is absent (yellow arrow) in the supraorbital mesenchyme of *Msx2-Cre Wls^fl/fl^* embryos at E11.5. (E) *Twist1* is expressed in the supraorbital mesenchyme of control embryos at E11.5. (F) Expression of *Twist1* is lost in the supraorbital mesenchyme of *Msx2-Cre Wls^fl/fl^* embryos at E11.5 (white arrow). (G and H) Expression of epithelial *Wnt3* is similar between control and *Msx2-Cre Wls^fl/fl^*embryos at E11.5. (I and J) Epithelial *Wnt4* levels are comparable between control and *Msx2-Cre Wls^fl/fl^* embryos at E11.5. (K and L) TUNEL assay indicates that levels of apoptosis are similar in control and *Msx2-Cre Wls^fl/fl^* embryos at E11.5. (M and N) Ki-67 staining at E11.5 shows that proliferation is reduced upon deletion of epithelial *Wls*. (O) Statistical analysis of Ki-67+ cells in supraorbital epithelium and underlying mesenchyme. Three pairs of control and mutant embryos were analyzed. Unpaired student *t*-test was used to evaluate significance. p<0.05 was considered significant. Scale bars: 50μm. The white dashed line indicates the boundary between epithelium and mesenchyme.

### Expression of *Fgf10* in eyelid mesenchyme requires epithelial Wnt **ligand secretion**

We observed that loss of mesenchymal β-catenin in *Prx1-Cre* β*-catenin^fl/fl^*mutants caused reduced proliferation in the overlying epithelium as well as in supraorbital mesenchyme, suggesting that canonical Wnt signaling in the mesenchyme controls expression of secreted signals required for epithelial proliferation. Previous studies showed that FGF10, which is expressed in eyelid mesenchyme, controls epithelial proliferation by binding to its receptor FGFR2 in the overlying epithelium [11]. Since *Fgf10* is a direct target gene of canonical Wnt signaling in other developmental contexts, we asked whether *Fgf10* expression was affected in the eyelid of *Wls-*deficient mutants. Because significant *Fgf10* expression in the upper eyelid initiates from E12.5 [11], and the upper eyelid of *Msx2-Cre Wls^fl/fl^* mutants fail to form, we utilized *Msx2- Cre^low^* mice that had been maintained for many generations and exhibited partial silencing of Cre expression. *Msx2-Cre^low^* mice exhibited periocular Cre activity that was not as broad as that of typical *Msx2*-*Cre* in supraorbital epithelium at E11.5, being more restricted to the eyelid groove (Fig. S3B-D). At E13.5, Cre activity in *Msx2-Cre^low^* mice was mainly present in the epithelium of the posterior part of the developing upper eyelid (Fig. S3E-G). *Msx2-Cre^low^ Wls^fl/fl^* mutants displayed an EOB phenotype (Fig. S4A and S4B), and H&E staining of sectioned tissue showed that the upper eyelids of the mutants formed but were hypoplastic at E18.5 (Fig. S4C and S4D). Analysis of upper eyelid development at E15.5 and E12.5 confirmed that the upper eyelids of *Msx2-Cre^low^ Wls^fl/fl^* mutants were significantly reduced in size compared to controls (Fig. S4E-H). Expression of LEF1 and proliferation were reduced in the hypoplastic upper eyelids of the mutants (Fig. 5A-D and 5G), similar to observations in *Prx1-Cre* β*-catenin^fl/fl^* and *Msx2-Cre Wls^fl/fl^* mutants. We found that *Fgf10* expression was lost in the upper eyelid of *Msx2-Cre^low^ Wls^fl/fl^* mutants (Fig. 5E and 5F). Together, these data demonstrate that epithelial Wnts are required for *Fgf10* expression in upper eyelid mesenchyme.

**Figure 5.**
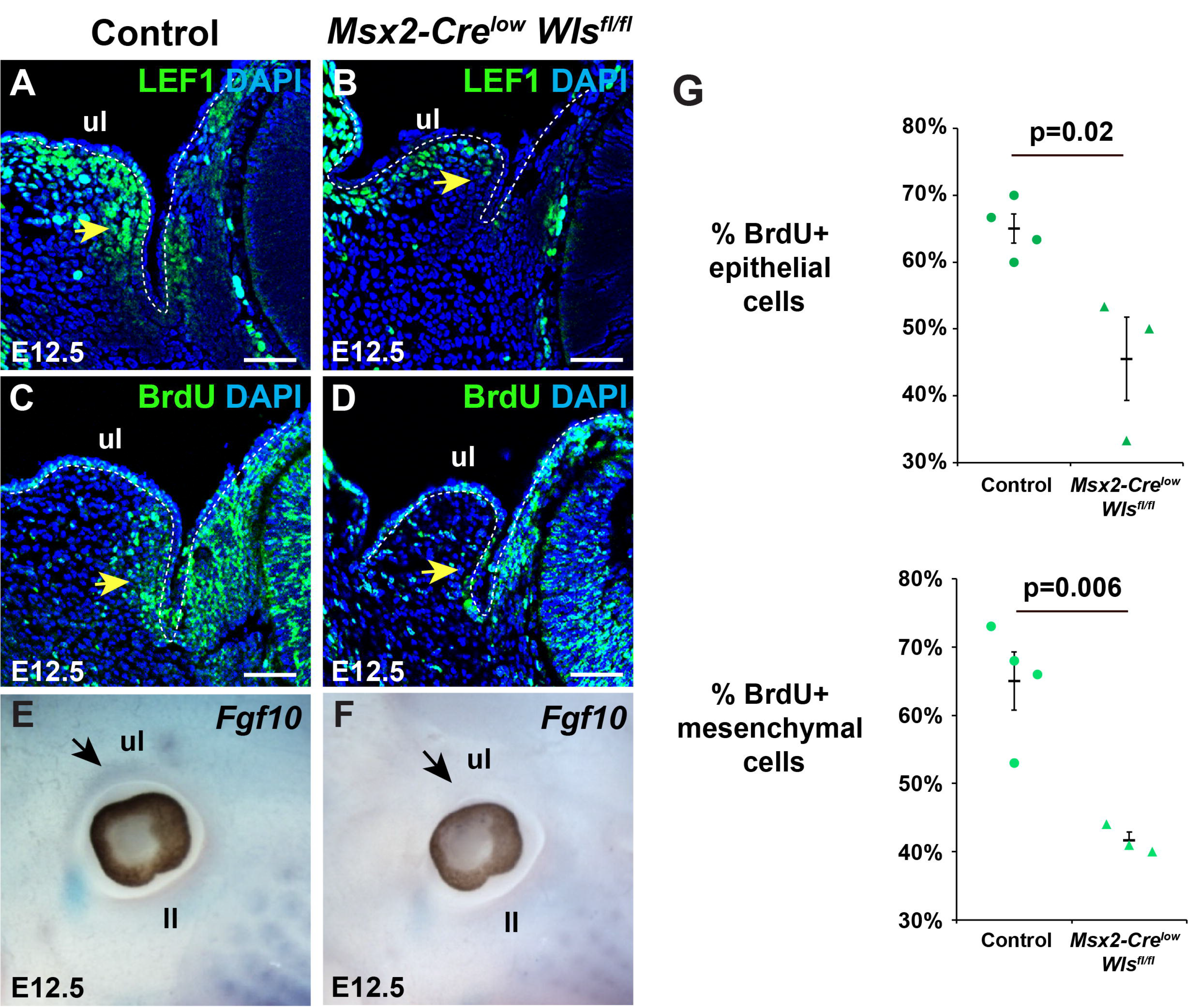
Expression of *Fgf10* in eyelid mesenchyme is dependent on secretion of epithelial Wnt ligands. (A) IF staining shows that LEF1 is strongly expressed in the mesenchyme of developing upper eyelid primordia of control embryos at E12.5 (yellow arrow). (B) IF staining shows that LEF1 expression is significantly reduced in the supraorbital mesenchyme of the hypoplastic upper eyelid primordia of *Msx2-Cre^low^ Wls^fl/fl^* embryos at E12.5 (yellow arrow). (C) BrdU staining shows that cell proliferation is enriched in the epithelium and posterior mesenchyme of upper eyelid primordia in control embryos at E12.5 (yellow arrow). (D) BrdU staining shows that proliferation is reduced in the upper eyelid primordia of *Msx2-Cre^low^ Wls^fl/fl^* embryos at E12.5 (yellow arrow). (E) *Fgf10* is expressed in the developing upper eyelid primordia of control embryos at E12.5 (black arrow). (F) *Fgf10* expression is undetectable in the developing upper eyelid primordia of *Msx2-Cre^low^ Wls^fl/fl^* embryos at E12.5 (black arrow). (G) Statistical analysis of BrdU+ cells in supraorbital epithelium and underlying mesenchyme. Four control and three mutant embryos were analyzed. Unpaired student *t*-test was used to evaluate significance. p<0.05 was considered significant. Scale bars: 50μm. The white dashed line indicates the boundary between epithelium and mesenchyme. Abbreviations: ul, upper eyelid; ll, lower eyelid.

### Expression of epithelial Wnts is regulated by p63 during eyelid development

In developing dorsal skin, epithelial p63 regulates transcription of multiple Wnt genes [26]. Since *p63* knockout mice also show an EOB phenotype [27], we asked whether p63 is required for Wnt expression in eyelid epithelium. We first examined the eyelids of *p63^+/-^* and *p63^-/-^* embryos. We found that heterozygous deletion of *p63* did not affect eyelid development. By contrast, the eyelids of homozygous *p63* knockout mutants were significantly smaller than those of wild-type or heterozygous mutant littermates at E14.5 and E17.5 (Fig. S5A-D). Expression of the p63 target genes KRT5 and KRT14 was reduced in developing eyelid epithelium of *p63^-/-^* embryos at E14.5 (Fig. 6A-D). We found rthar expression of the Wnt ligands *Wnt3* and *Wnt4* was also down- regulated in *p63*-deleted eyelid epithelium (Fig. 6E-H). In line with this, expression of the Wnt target genes *Lef1* and *Twist1* was also decreased in *p63^-/-^* mutant eyelids (Fig. 6I-L). Taken together, these data indicate that p63 regulates expression of epithelial Wnt genes during eyelid development.

**Figure 6.**
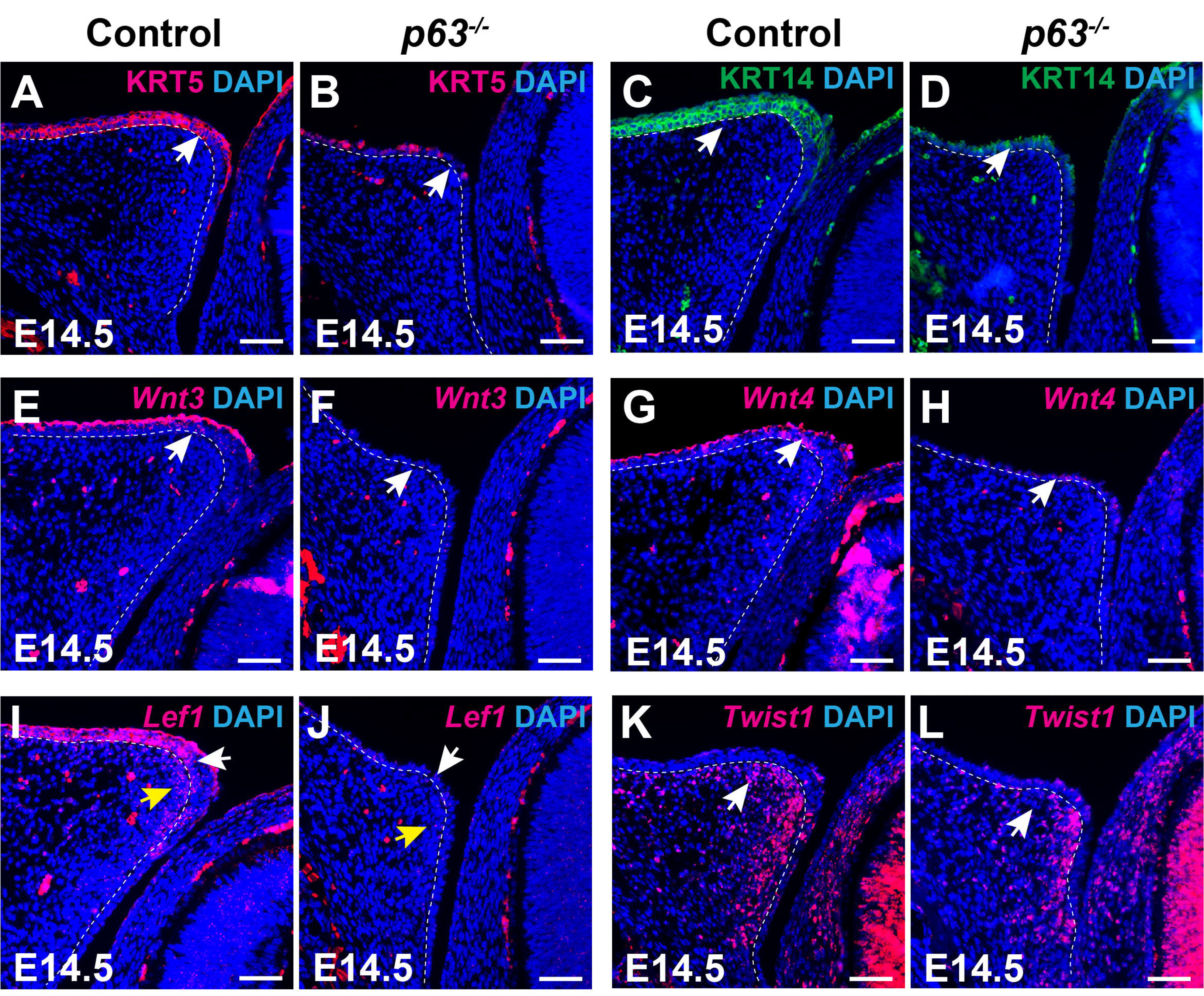
p63 regulates expression of Wnt genes in eyelid epithelium. (A and C) KRT5 and KRT14 are expressed in the epithelium of the upper eyelid of control embryos at E14.5 (white arrows). (B and D) KRT5 and KRT14 expression is diminished in the epithelium of the upper eyelid of E14.5 *p63^-/-^*embryos (white arrows). (E and G) *Wnt3* and *Wnt4* are expressed in the epithelium of the upper eyelid of control embryos at E14.5 (white arrows). (F and H) Expression of *Wnt3* and *Wnt4* is significantly reduced in the epithelium of the upper eyelid of *p63^-/-^* embryos at E14.5 (white arrows). (I) *Lef1* is expressed in the epithelium (white arrow) and underlying mesenchyme (yellow arrow) of the upper eyelid of control embryos at E14.5. (J) Expression of *Lef1* is lost in both epithelium (white arrow) and underlying mesenchyme (yellow arrow) of the upper eyelid of E14.5 *p63^-/-^* embryos. (K) *Twist1* is expressed in the mesenchyme of the upper eyelid of control embryos at E14.5 (white arrow). (L) Expression of *Twist1* is reduced in the mesenchyme of the upper eyelid of E14.5 *p63^-/-^* embryos. Scale bars: 50μm. The white dashed line indicates the boundary between epithelium and mesenchyme.

## Discussion

Reciprocal interactions between epithelium and underlying mesenchyme are critical for the development of multiple organs and appendages, including skin, lung, limb and eyelid [3, 25, 28, 29]. Eyelid development is a multi-stage process that is controlled by distinct networks involving diverse signaling pathways and transcription factors. Among these, FGF10 secreted from eyelid mesenchyme binds to its receptor FGFR2b, which is expressed in the overlying epithelium, to modulate proliferation and migration of epithelial cells [11]. This indicates the existence of mesenchyme-to-epithelium signaling during eyelid development. By comparison, the signaling relays from epithelium to mesenchyme required for eyelid formation are less well characterized.

Wnt ligands are secreted factors that mediate epithelial-mesenchymal interactions in various contexts. As examples, ectodermal Wnt ligands are required to maintain the outgrowth of mesenchyme during limb development [20]; and epithelial Wnt secretion regulates proliferation and differentiation of mesenchymal progenitors in the skin dermis [25] and in the developing lung [30]. During eyelid development, multiple Wnt genes including *Wnt3* and *Wnt4* show persistent expression in periocular epithelium and Wnt receptors such as FZD7 are expressed in the underlying mesenchyme [16]. These observations suggested that epithelial Wnt ligands might play key roles in mediating epithelial-mesenchymal interactions during eyelid development.

In support of this hypothesis, we found that deletion of mesenchymal β- catenin or prevention of epithelial Wnt ligand secretion resulted in failure of upper eyelid development, indicating that epithelial Wnts regulate eyelid development by activating canonical Wnt signaling in the mesenchyme. We further uncovered that deletion of β-catenin in developing eyelid mesenchyme abrogates *Fgf10* expression, consistent with prior data identifying *Fgf10* as a Wnt/β-catenin target gene [31]. Together with prior findings that *Fgf10* knockout mice display reduced proliferation of eyelid epithelium [11], our data suggest a model in which epithelial Wnt ligands signal to the mesenchyme to activate *Fgf10* expression, which then signals back to the epithelium to control its proliferation. Similar regulation of mesenchymal *Fgf10* by epithelial Wnts is observed in other developing organs, such as limb and lung [20, 30], indicating that the Wnt-FGF10 axis is a common mechanism underlying epithelial-mesenchymal interactions. While *Fgf10* mutants display reduced proliferation only in the epithelium of the eyelid [11], we find that canonical Wnt signaling regulates cell proliferation in presumptive eyelid mesenchyme as well as in the epithelium. These observations indicate that the Wnt/β-catenin pathway controls eyelid development by additional mechanisms.

Overactivation of Wnt/β-catenin signaling is also known to disrupt eyelid development. For example, genetic deletion of the secreted Wnt inhibitor DKK2 or constitutive activation of epithelial β-catenin both cause EOB phenotypes [18, 32]. Therefore, it is crucial to precisely control activity of epithelial Wnt ligands during eyelid development. However, the mechanisms that regulate their expression have been unclear. To address this question, we examined a possible role for the epithelial transcription factor p63 in regulating epithelial Wnt gene expression.

p63 is a master regulator of epidermal development [33], and genetic deletion of *p63* results the failed development of multiple organs and tissues including the epidermis, limb and eyelid [27]. p63 controls epidermal differentiation by orchestrating the expression and activity of multiple transcription factors and signaling pathways. In the developing skin epidermis, p63 directly activates transcription of epidermal Wnt genes to regulate epidermal differentiation [26]. Consistent with this observation, we find that homozygous deletion of *p63* impairs expression of several epithelial Wnt genes and causes reduced levels of canonical Wnt signaling in the developing eyelid. However, it is noteworthy that hypoplastic eyelids still develop in homozygous *p63* mutants, suggesting that additional mechanisms contribute to the regulation of epithelial Wnt gene expression.

In summary, our data provide genetic evidence that Wnt/β-catenin signaling controls eyelid development by mediating epithelial-mesenchymal interactions. These findings provide a mechanistic basis for investigating the etiology of eyelid colobomas.

## Acknowledgments

This work was supported in part by NIAMS/NIH grant R37AR047709 (SEM) and the National Nature Science Foundation of China (31970775) (GM)

## Author contributions

Conceptualization: XZ; Formal Analysis: XZ; Funding Acquisition: SEM, GM; Investigation: XZ; Methodology: XZ; Project Administration, SEM, GM; Resources: MS, SEM, GM; Supervision: SEM, GM; Validation: XZ; Visualization: XZ; Writing - Original Draft Preparation: XZ; Writing - Review and Editing: XZ, SEM, MG.

## Data availability statement

The authors declare that the main data supporting the findings of this study are available within the article and its Supplemental Information files. All correspondence and material requests should be addressed to Sarah E. Millar or Gang Ma. This study includes no data deposited in external repositories.

---

Figure S1. ***Prx1-Cre* deleted cells are present in supraorbital and developing upper eyelid mesenchyme.** (A) Schematic of the lateral view of the head of a mouse embryo. (B-J) Tissue (B,E,H) and tissue sections (C,D,F,G,I,J) from *Prx1-Cre* embryos carrying the *ROSA26R^mT/mG^* Cre reporter allele. (B) GFP, indicating Cre reporter expression, is present in a subset of cranial facial mesenchymal cells at E11.5. (C) Frontal section corresponding to the position indicated by the yellow dashed line in (B); GFP+ cells are present in supraorbital mesenchyme. (D) Amplified view of the area outlined by the yellow dashed rectangle in (C). (E) GFP+ cells in periocular mesenchyme expand rostrally at E12.5. (F) Frontal section corresponding to the position indicated by the yellow dashed line in (E); GFP+ cells are broadly present in developing upper eyelid mesenchyme. (G) Amplified view of the area outlined by the yellow dashed rectangle in (F). (H) GFP+ cells in periocular mesenchyme expand further rostrally at E13.5. (I) Frontal section corresponding to the position indicated by the yellow dashed line in (H); GFP+ cells are uniformly present in the developing upper eyelid mesenchyme. (J) Amplified view of the area outlined by the yellow dashed rectangle in (I). Scale bars: 500μm in panels (B, E and H); 100 μm in panels (C, D, F, G, I and J). The white dashed line in (D,G,J) indicates the boundary between epithelium and mesenchyme. Abbreviations: ul, upper eyelid; ll, lower eyelid.

Figure S2. ***Msx2-Cre* deleted cells are present in periocular epithelium of the developing eyelid.** (A) Schematic of the lateral view of the head of a mouse embryo. (B-J) Tissue (B,E,H) and tissue sections (C,D,F,G,I,J) from *Msx2-Cre* embryos carrying the *ROSA26R^mT/mG^* Cre reporter allele. (B) GFP, indicating Cre reporter expression, is broadly present in head epithelium at E11.5. (C) Frontal section corresponding to the position indicated by the yellow dashed line in (B); GFP+ cells are present in supraorbital epithelium. (D) Amplified view of the area outlined by the yellow dashed rectangle in (C). (E) GFP+ cells in the periocular epithelium at E12.5. (F) Frontal section corresponding to the position indicated by the yellow dashed line in (E); GFP+ cells are broadly present in the developing upper eyelid epithelium. (G) Amplified view of the area outlined by the yellow dashed rectangle in (F). (H) GFP+ cells are present in periocular epithelium at E13.5. (I) Frontal section corresponding to the position indicated by the yellow dashed line in (H); GFP+ cells are present in the developing upper and lower eyelid epithelium. (J) Amplified view of the area outlined by the yellow dashed rectangle in (I). Scale bars: 500μm in panels (B, E and H), 100 μm in panels (C, D, F, G, I and J). The white dashed line in (D,G,J) indicates the boundary between epithelium and mesenchyme. Abbreviations: ul, upper eyelid; ll, lower eyelid.

Figure S3. ***Msx2-Cre^low^* directs deletion in epithelium of the developing eyelid groove and eyelids.** (A) Scheme of the lateral view of the head of a mouse embryo. (B-G) Tissue whole mounts (D,E) and sections (C,D,F,G) from *Msx2-Cre^low^* embryos carrying the *ROSA26R^lacZ^* Cre reporter allele. (B) LacZ+ cells are present in the epithelium of the eyelid groove at E11.5. (C) Frontal section corresponding to the position indicated by the black dashed line in (B); LacZ+ cells are present in the epithelium of the eyelid groove. (D) Amplified view of the area outlined by the black dashed rectangle in (C). LacZ+ cells are present in the epithelium of the eyelid groove (black arrowhead), and sporadic and weak LacZ+ cells can be identified in the supraorbital epithelium (red arrowhead). (E) LacZ+ cells are present in the epithelium of the eyelid at E13.5. (F) Frontal section corresponding to the position indicated by the black dashed line in (E); LacZ+ cells are present in the developing eyelid epithelium. (G) Amplified view of the area outlined by the black dashed rectangle in (F). LacZ+ cells are enriched in the epithelium of posterior upper eyelid (black arrowheads). Abbreviations: ul, upper eyelid; ll, lower eyelid.

Figure S4. ***Msx2-Cre^low^ Wls^fl/fl^* mutants exhibit hypoplastic eyelids.** (A) The eyelids of control pups are fused at P0. (B) The eyes of *Msx2-Cre^low^ Wls^fl/fl^* pups exhibit EOB at P0. (C) H&E staining shows that eyelids are fused in control embryos at E18.5. (D) H&E staining indicates that the upper eyelid is hypoplastic in *Msx2-Cre^low^ Wls^fl/fl^* embryos (black arrow). (E) H&E staining shows that control eyelids are well-formed at E15.5. (F) H&E staining indicates that the upper eyelid is significantly smaller in *Msx2-Cre^low^ Wls^fl/fl^* embryos at E15.5 compared with controls (black arrow). (G) H&E staining shows that at E12.5, control embryos have morphologically distinguishable eyelid primordia. (H) H&E staining shows that in *Msx2-Cre^low^ Wls^fl/fl^* embryos, the size of upper eyelid primordia is significantly reduced at E12.5 compared with controls (black arrow). Abbreviations: ul, upper eyelid; ll, lower eyelid.

Figure S5. ***p63^-l-^* mutants exhibit hypoplastic eyelids.** (A) H&E staining shows that the eyelids of control embryos are fused at E17.5. (B) H&E staining shows that the eyes of *p63^-l-^* embryos are open at E17.5. (C) H&E staining shows that eyelids are well-formed in control embryos at E14.5. (D) H&E staining indicates that eyelids are hypoplastic in *p63^-l-^* embryos at E14.5. Scale bars: 150μm. Abbreviations: ul, upper eyelid; ll, lower eyelid.

## Notes

### Competing Interest Statement

The authors have declared no competing interest.

