## Supplementary figures and images for "Wnt/β-catenin signaling controls mouse eyelid growth by mediating epithelial-mesenchymal interactions"

### Fig.S1

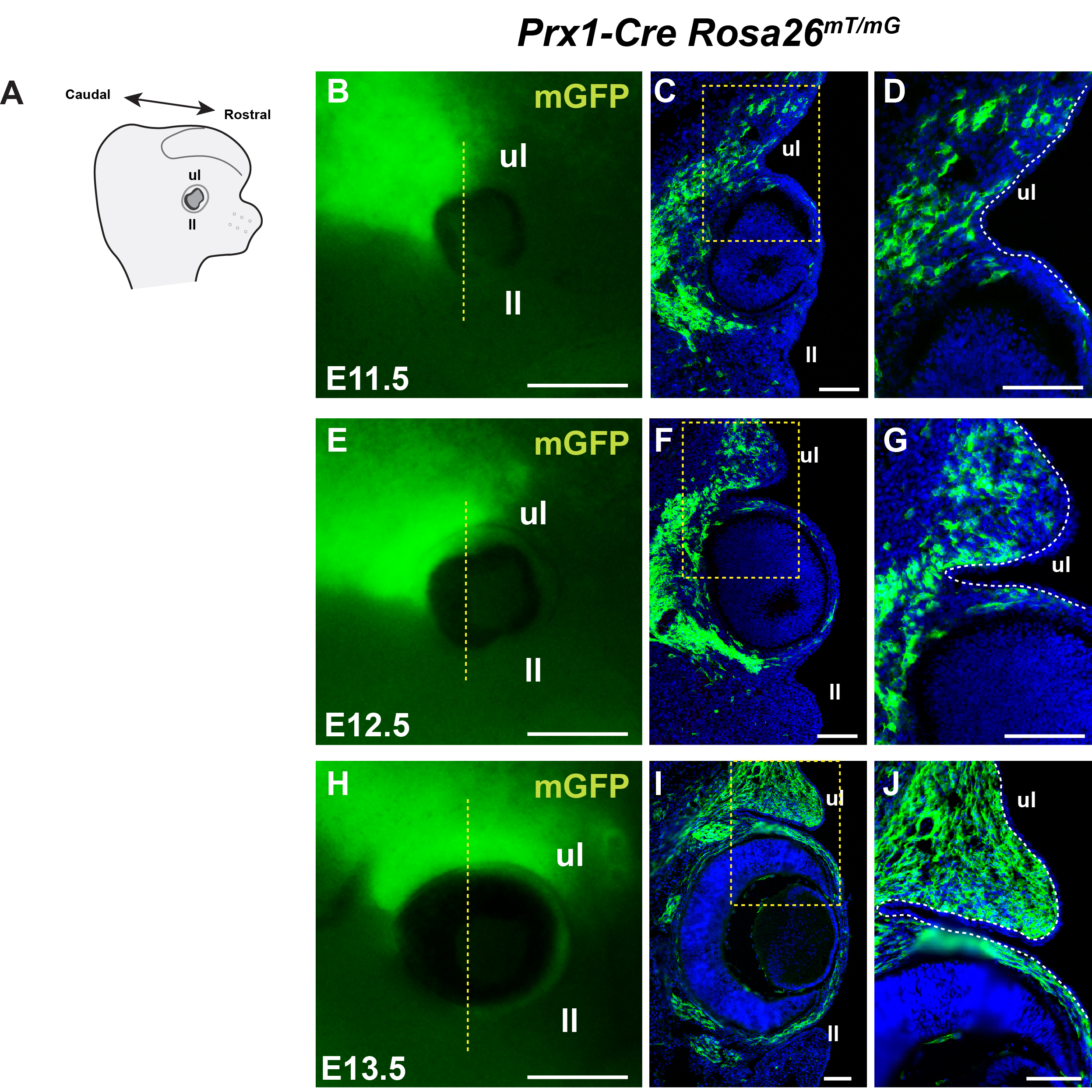

### Fig.S2

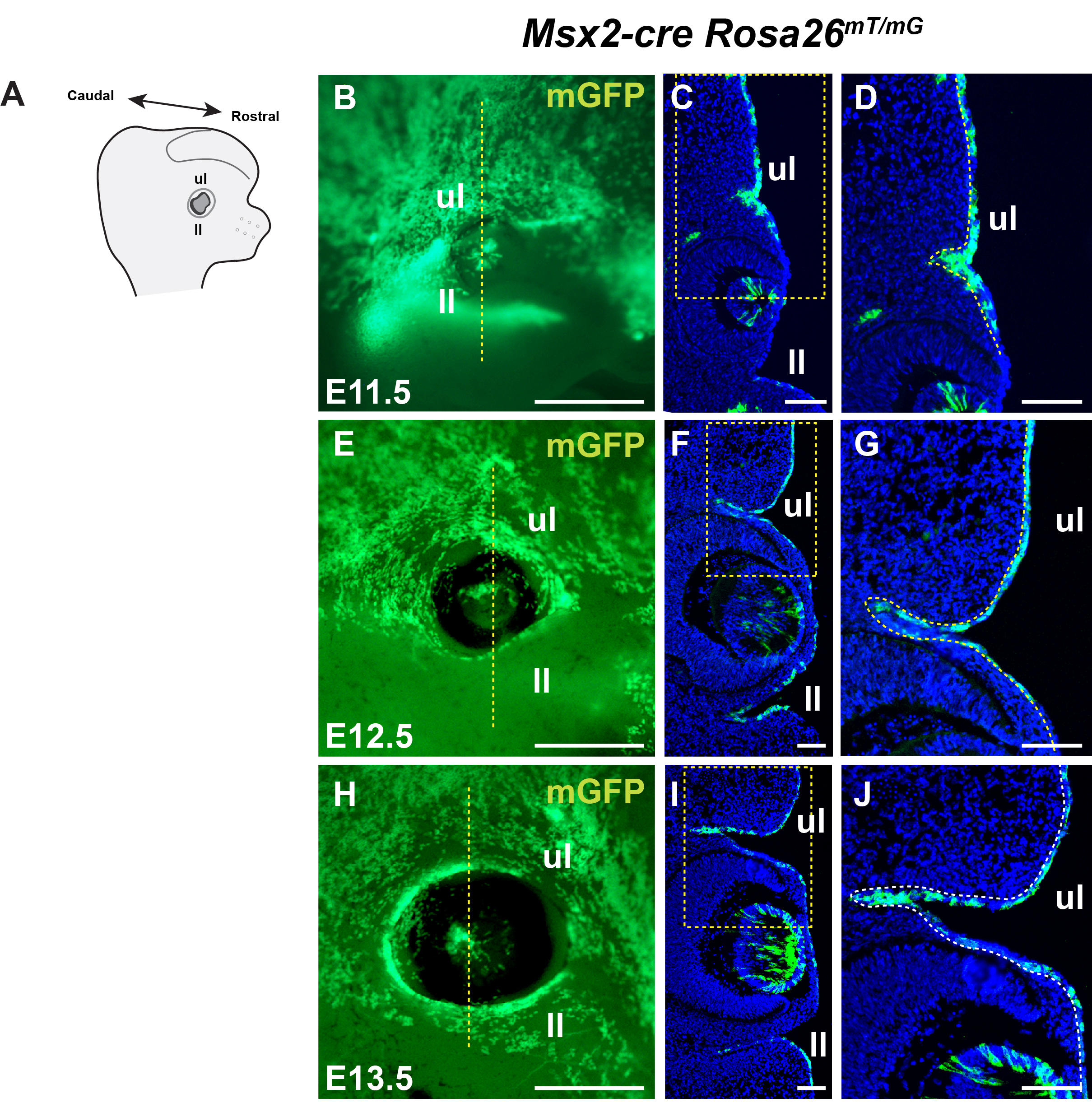

### Fig.S3

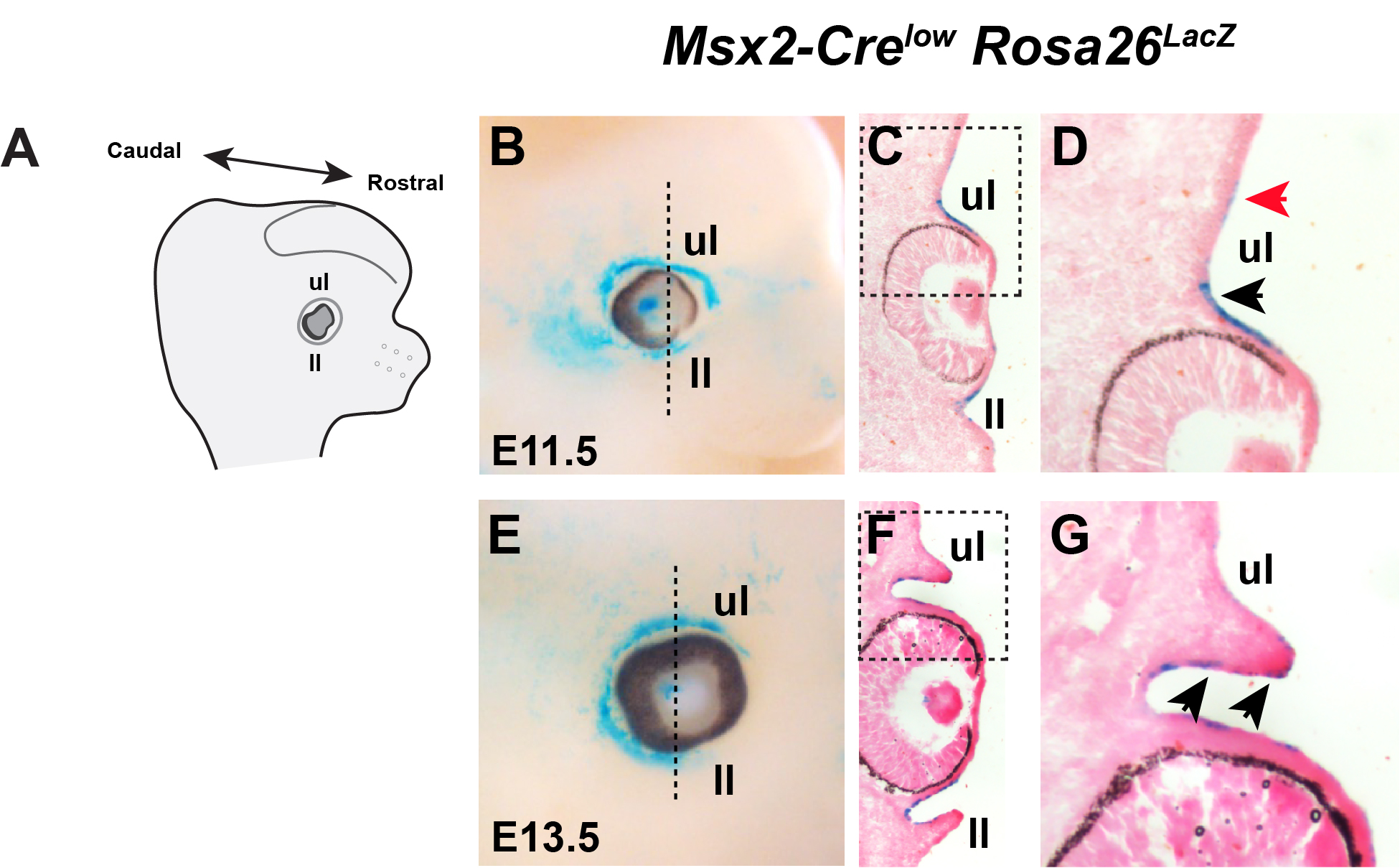

### Fig.S4

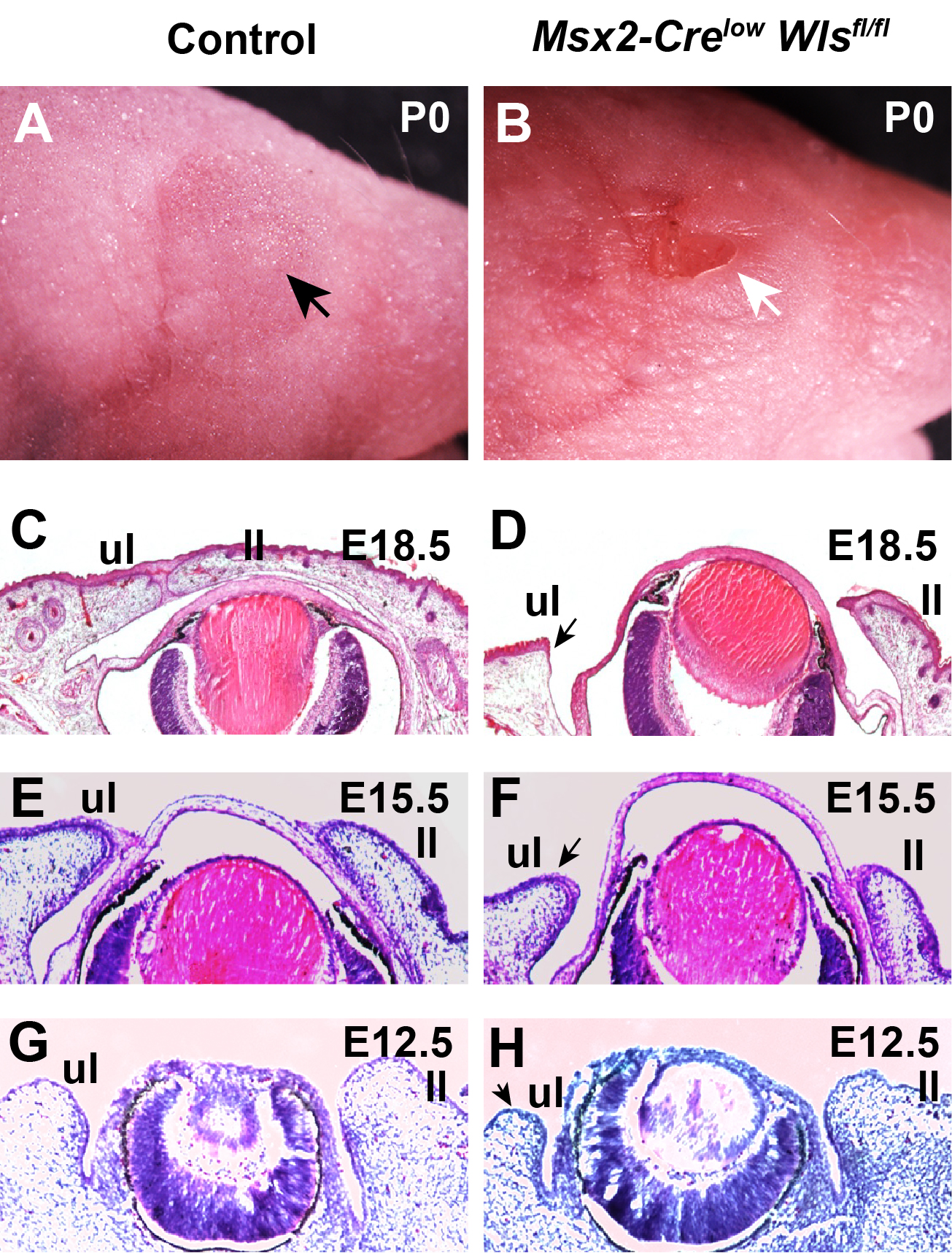

### Fig.S5

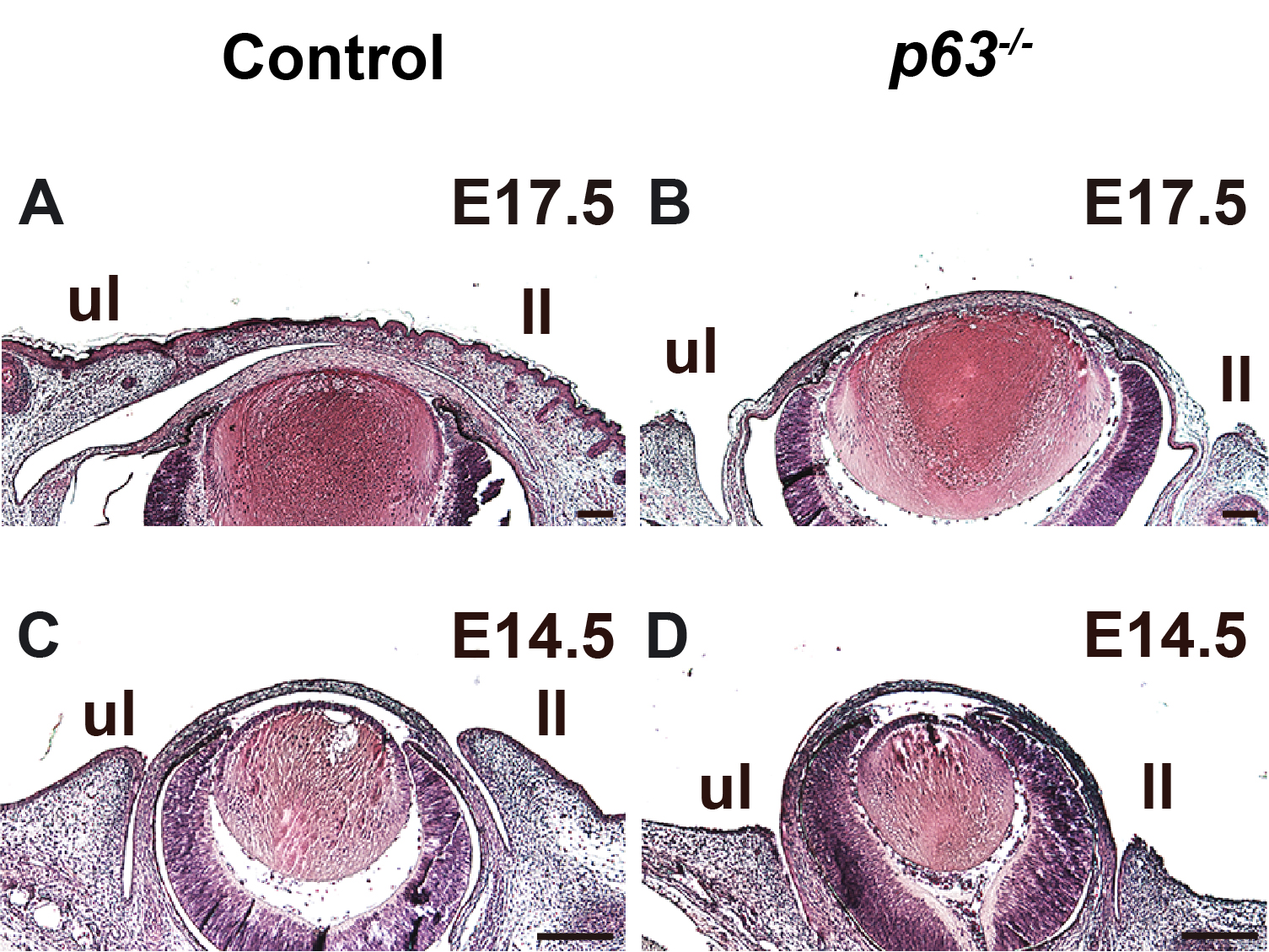
